# Multiple SARS-CoV-2 introductions shaped the early outbreak in Central Eastern Europe: comparing Hungarian data to a worldwide sequence data-matrix

**DOI:** 10.1101/2020.05.06.080119

**Authors:** Gábor Kemenesi, Safia Zeghbib, Balázs A Somogyi, Gábor Endre Tóth, Krisztián Bányai, Norbert Solymosi, Peter M Szabo, István Szabó, Ádám Bálint, Péter Urbán, Róbert Herczeg, Attila Gyenesei, Ágnes Nagy, Csaba István Pereszlényi, Gergely Csaba Babinszky, Gábor Dudás, Gabriella Terhes, Viktor Zöldi, Róbert Lovas, Szabolcs Tenczer, László Kornya, Ferenc Jakab

**Affiliations:** Virological Research Group BSL-4 Laboratory, Szentágothai Research Centre, University of Pécs, Pécs, Hungary; Institute of Biology, Faculty of Sciences, University of Pécs, Pécs, Hungary; National Coronavirus Research Centre, University of Pécs, Pécs, Hungary; Institute for Veterinary Medical Research, Centre for Agricultural Research, Budapest, Hungary; University of Veterinary Medicine Budapest, Centre for Bioinformatics, Budapest, Hungary; Translational Discovery, Stromal Biology, Bristol-Myers Squibb, Princeton, NJ, USA; National Food Safety Office, Budapest, Hungary; Szentágothai Research Centre, Bioinformatics Research Group, Genomics and Bioinformatics Core Facility, University of Pécs, Pécs, Hungary; Clinical Research Centre, Medical University of Bialystok, Bialystok, Poland; Hungarian Defense Forces, Military Medical Centre, Budapest, Hungary; Institute of Clinical Microbiology, University of Szeged, Faculty of Medicine, Szeged, Hungary; Independent researcher, Vantaa, Finland; Institute for Computer Science and Control, Hungarian Academy of Sciences; Central Hospital of Southern Pest, Budapest

## Abstract

Severe Acute Respiratory Syndrome Coronavirus 2 is the third highly pathogenic human coronavirus in history. Since the emergence in Hubei province, China, during late 2019 the situation evolved to pandemic level. Following China, Europe was the second epicenter of the pandemic. To better comprehend the detailed founder mechanisms of the epidemic evolution in Central-Eastern Europe, particularly in Hungary, we determined the full-length SARS-CoV-2 genomes from 32 clinical samples collected from laboratory confirmed COVID-19 patients over the first month of disease in Hungary. We applied a haplotype network analysis on all available complete genomic sequences of SARS-CoV-2 from GISAID database as of the 21th of April, 2020. We performed additional phylogenetic and phylogeographic analyses to achieve the recognition of multiple and parallel introductory events into our region. Here we present a publicly available network imaging of the worldwide haplotype relations of SARS-CoV-2 sequences and conclude the founder mechanisms of the outbreak in Central-Eastern Europe.

## Introduction

Following the 2002 SARS (Severe Acute Respiratory Syndrome) pandemic and the discovery of MERS (Middle Eastern Respiratory Syndrome) coronavirus in 2012, the third highly pathogenic human coronavirus in history emerged in Hubei province, China, during late 2019. The novel virus was subsequently named Severe Acute Respiratory Syndrome Coronavirus 2 (SARS-CoV-2) and the acute respiratory disease as coronavirus disease 19 (COVID-19) ^1^. Currently, SARS-CoV-2 is responsible for the ongoing coronavirus pandemic spreading on all inhabited continents. As of April 26, 2020, the confirmed case numbers surpassed 3 million worldwide and the disease associated mortality rate exceeded 200,000 ^2^.

At the onset of the second week of March, Europe became the next epicenter of the pandemic, following China, as reported by the World Health Organization ^3^. By the end of April, more than one million laboratory confirmed cases were reported from all European countries ^4^. The first two Hungarian cases were officially confirmed on March 4^th^, according to the data of ECDC Communicable Disease Threats Report ^5^. Border closures and universal ban regarding public gatherings was announced on March 17.

To better comprehend the detailed founder mechanisms of the epidemic evolution in Central-Eastern Europe, particularly in Hungary, we determined the full-length SARS-CoV-2 genomes from 32 clinical samples collected from laboratory confirmed COVID-19 patients over the first month of disease in Hungary. Our virus sampling started from this date and spanned the first two weeks of country-wide mitigation regulations (March 17 through April 2, 2020).

## Results and Discussion

In order to understand the origin of Hungarian-based COVID-19 epidemics and provide baseline data for the evaluation of future epidemic events we applied a network analysis on complete genomic sequence data of SARS-CoV-2 available in GISAID database^6^ current to April 21. The network showed negative exponential degree distribution which is common regarding scale-free networks^7^. This characteristic network is typical for epidemics^8^. However, several nodes represented a higher frequency in the lower part of the plot which is the tendency associated with small-world networks^9^ (Supplementary Figure 1.). Altogether, a total of 147 clusters were identified with a Girvan-Newmann community detection algorithm^10^. In consideration of this approach, a total of nine main clusters were described which together serve as the base for the remaining smaller clusters (**Figure 1**.). Although the investigated network contained relatively high number of clusters, its diameter is 25, which infers the farthest distance in the matrix between two sequence is 25 steps, whilst the average path length is 8.91 steps. The high cluster rate was supported by the ratio of these two measures. The proportion of present edges from all possible edges in the network was 0.004 (edge density).

**Figure 1.**
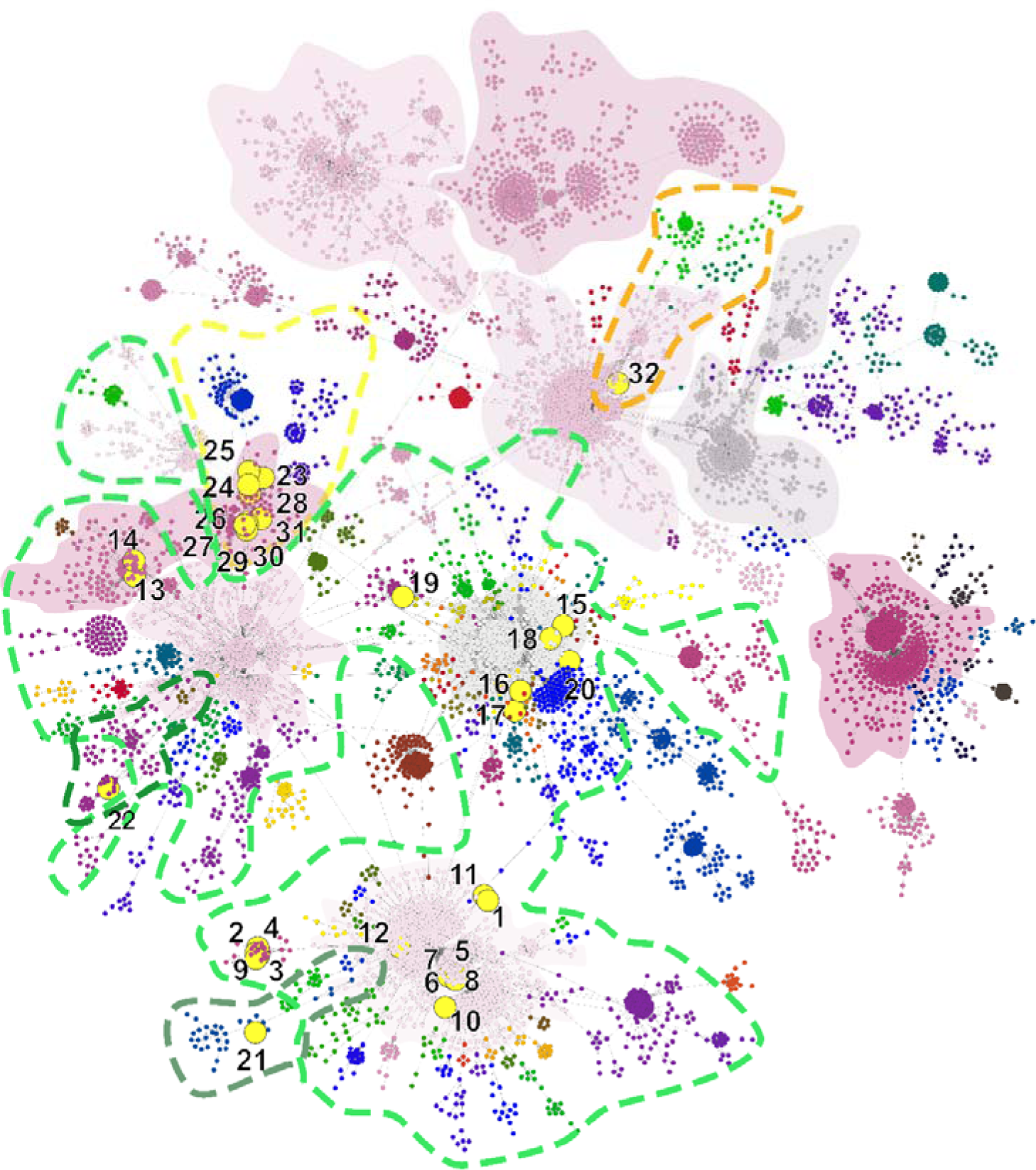
Genetic network analysis of 7864 SARS-CoV-2 complete genomic sequences. Hungarian strains are indicated with numbered yellow dots – numbers referring to Table 1. The nine major clades are represented by a solid color. Genetic lineages are marked with colored dotted lines, where green lines are bordering B 1, B 1.1 and B 1.11; yellow and orange lines mark B 1.5 and B 3 respectively.

Hungarian genomes are dispersed within four main clusters out of the nine (**Figure 2.**). The genome designated SARS-CoV-2/human/Hungary/620/27_03_2020 is positioned in the main Chinese cluster (incorporating early Chinese sequences and data from Hangzhou, in January) and closely connected to a Taiwanese sequence on March 23rd (**Figure 2**., Cluster C). Apart from other Hungarian sequences, this is the only indication for the introduction from a non-European source towards Hungary at the examined time-period in consideration of the available sequence data. The remainder of the sequences are dispersed among three other main clusters: A, B and I (**Figure 2**.). A and I clusters are structured by mostly the Western-European sequences, whilst B is a dominant cluster in the USA. Although sampling bias may largely alter the conclusions for the exact geographic origin of a particular strain, the main patterns as multiple introductions from different sources can be concluded.

**Figure 2:**
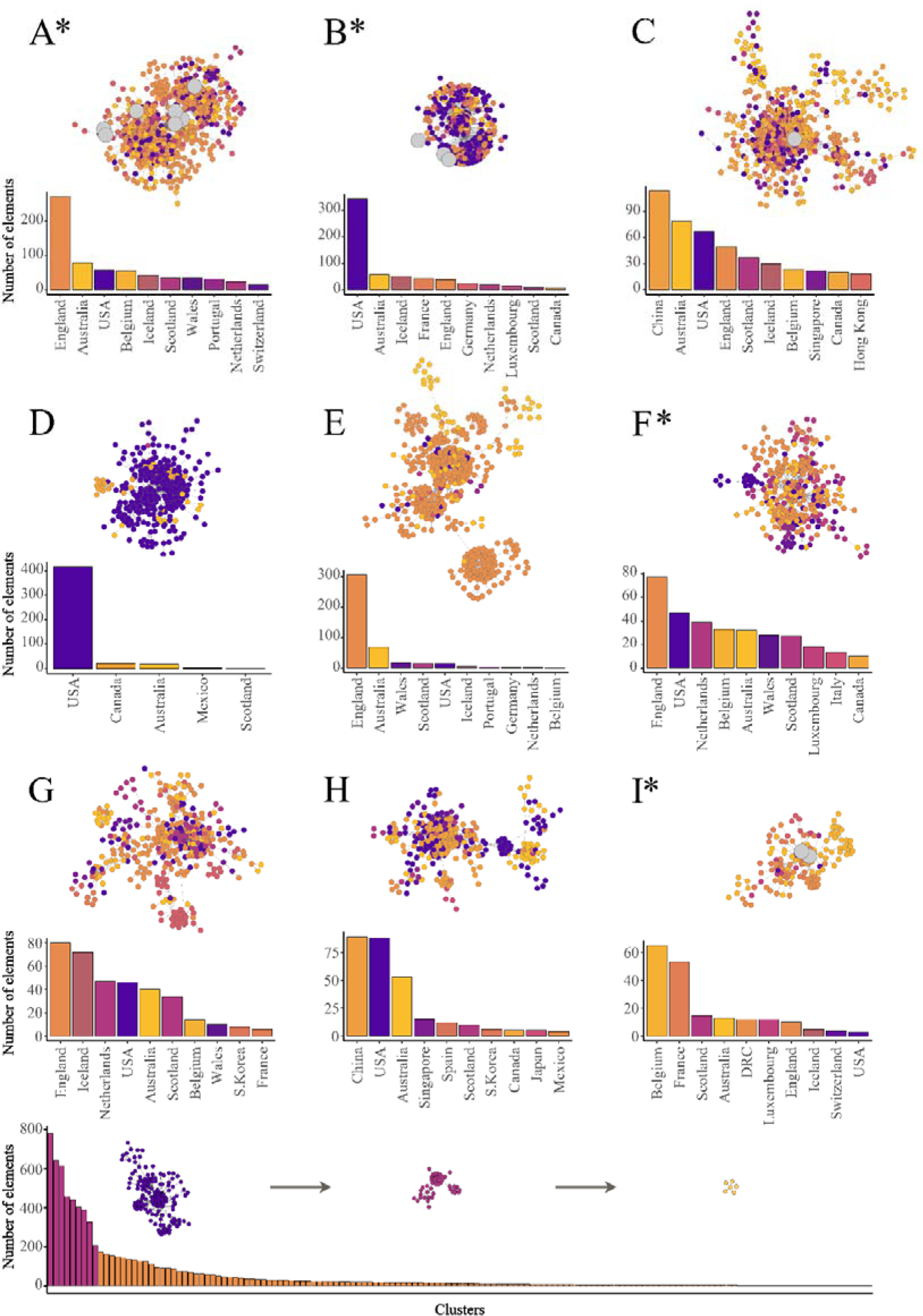
Representation of the nine main clusters which serve as the baseline regarding the worldwide haplotype network of SARS-CoV-2 genomes as of April 21, 2020. The ten most common countries of each cluster are summarized in a column graph and represented using different colors. Hungarian sequences are depicted by enlarged grey dots. Star indicates the participation of a sample from this study indirectly to the specified main cluster (i.e., participation in a sub-cluster relevant to the main cluster). Number of elements within each remaining (n=147) smaller cluster is indicated as a simple column chart at the bottom of the figure.

Using the complete haplotype network dataset as a backbone, we applied additional phylogenetic and phylogeographic analyses (Supplementary Figure S2). It is likely that occupation-related movement within the EU resulted in multiple introductory events from Western-European host countries towards Central-Eastern Europe. This observation is further supported by recent narrative analysis on the Nextstrain online platform focusing on Eastern European processes of SARS-CoV-2 pandemic evolution^11^. Similarly to Hungary and possibly to the entire region, there were eleven separate introductions to Poland, based on the currently available sequence data^11^. In order to leverage additional support regarding this phenomenon, we applied a local Nextstrain database workflow in the addition of the sequences from this manuscript (Supplementary Figure S3)^12^. As a result of this analysis and considering the observation from Poland, we were able to lend more support for the regular and dispersed introductions into Central-Europe. In addition to regular movement, the border restrictions as outbreak mitigation measures fixed a narrow timescale for individuals returning to Hungary and likely facilitated the parallel introductory events dispersed throughout the country. Based on genetic lineage categorization using PANGOLIN software, 20 out of the total 32 Hungarian sequences fell into the most dominant (i.e., most sequenced) lineage B.1 (**Table 1**). Dominance may largely depend on sampling heterogeneity between geographic regions and countries. However, it substantiates the connection of Hungary regarding SARS-CoV-2 cases to multiple European sources and provides additional support for the network analysis.

**Table 1**: Summary of the PANGOLIN software analysis. The table indicates the numbers of Figure 1. and offers additional details for each sample. Background data is also noted where it was available. Different background color of SARS-CoV-2 strain names highlight different genetic lineage groups represented with the same colored dotted lines on Figure 1

Across the phylogenetic tree (Supplementary Figure S2A), several of the Hungarian sequences were interspersed and mainly clustered with European sequences (England, France, Iceland and Germany) and supported with high posterior probabilities (>80%) while only one Hungarian sequence clustered with a North-American sequence (PP = 95%). These observations elegantly support the scenario regarding multiple individual introductions. In parallel, local clusters were also observed (PP = 100%) indicating local transmission even within the short timeframe of sampling. Moreover, several of the local clusters had very low PP indicating missing data which is likely to be the consequence of insufficient contact tracing and subsequent missing sequence data.

Within our dataset, the phylogeographic analysis indicated China as the root location (diffusion origin) (Supplementary Figure S2B). Moreover, the virus seemed to spread out to Hungary mainly from Western European countries, nevertheless local transmissions also contributed to disease spread within the country. The data correspond with the epidemiological history of SARS-2-CoV-2 in Hungary^4^.

As a main result of this study, we present and provide a large-scale haplotype network backbone in reference to the immediate analysis of pandemic evolution of SARS-CoV-2. It is a rapid and useful tool to assess the origin of particular sequences and the acquisition of important data for regarding public health mitigation actions, discovering unidentified infection sources or super-spreading events on a large-scale. In general, it provides the network-based opportunity of rapid, genetic distance-based analysis for all available sequence data, in any context. Herein, we offer this network file available for any researchers to facilitate the understanding of SARS-CoV-2 pandemic evolution. The network file is suitable to visualize any available sequences, available at late April 2020, in its context to all known sequence data.

The importance of early, country-based mitigation measures are thoroughly exemplified on this dataset. We presented the emergence of multiple virus clusters from various sources in Hungary during the early phase of the epidemic. However, the publicly available epidemiologic data indicate a predominance of confirmed cases in and adjacent to the capital city, Budapest. Possibly, this phenomenon is due to effective mitigation by limiting individual movement, application of social distancing and border restrictions^13^. Therefore, we believe a pan-European, coordinated mitigation policy will be beneficial to prevent significant mixture of European clusters during future epidemics.

Our research further highlights the importance of genomic epidemiologic tools for public health decision making. The combination of different methods (i.e., network analysis and phylogenetic approaches) may greatly facilitate the understanding of COVID-19 outbreak evolution.

## Methods

### Sample collection

Oro-pharyngeal swab samples were obtained from 32 patients during the period from March 17 to April 2. Within the frame of a country-wide collaboration network regarding SARS-CoV-2 research, nucleic-acid samples were received from University Hospitals at Szeged and Budapest and from the Hungarian Defense Forces, Military Medical Center. Ethical approval was obtained from the University of Pécs, Ethics Committee, under the registration number: 8218-PTE2020.

### Direct sequencing and primary data analysis from patient samples

Nucleic acid samples were extracted directly from oro-pharyngeal swab samples using a Direct-zol™ RNA MiniPrep Plus extraction kit (Zymo Research) and in full compliance to the manufacturers’ recommendations. Reverse transcription and multiplex PCR were performed on the basis of information provided by the Artic Network initiative ^14^. Both the concentration and the quality of the PCR products were measured and checked using the Agilent 4200 TapeStation System and ThermoFisher Scientific Qubit 3 Fluorometer. The 32 sequencing libraries were prepared using 98 overlapping amplicons covering the whole viral genome. The libraries were then quantitatively checked, barcoded and sequenced on 5 flow cells using Oxford Nanopore MinION Flow Cells (R9.4.1).

During primary data analysis, we used RAMPART to track the sequencing process in “real-time” in order to acquire instant information regarding the quality of samples and the coverage of the amplicons. Sequencing reads of samples with sufficient amplicon coverage were mapped and consensus sequences generated by the bioinformatics pipeline built within the Artic Network protocol.

### Genome data analysis

SARS-CoV-2 genomes (n=7864) were downloaded from GISAID database on 21 April, 2020. Only complete (>29000 base-pair length) and high quality (with <1% Ns, <0.05% unique amino acid mutations and no insertion/deletion unless verified by submitter) sequences were used for network construction. To quantify the sequence similarity, percent identity was calculated based on the BLAST ^15^ alignment for each paired sequence.

First, using the resulted similarity matrix, a fully connected, edge-weighted network was constructed, where each node represented a COVID sequence, while the edges represented their potential connections, and the edge weights (similarity values). Secondly, the edge weights were transformed (100-weight) in the full network to make high values low and low values high. Next, a minimum spanning tree (MST) was identified. In a spanning tree, every node has only one or two connections. If multiple edges have the same minimum weight, the algorithm will randomly pick one and not select all links with the same values. To manage this issue, the graph with additional edges was modified by adding every edge for each node having an equal or higher weight than the edges in the initial MST to the corresponding node. All data analyses were performed using the R 3.6.2 on Linux^16^, for network creation, and the Igraph package was applied^17^.

In regards to the generation of time-scaled phylogenetic tree, 105 SARS-CoV-2 genomes were retrieved from GISAID ^6^ following a manual selection based on the network analysis. The sequences were aligned in MAFFT ^18^ with default parameters. Subsequently, both best-fitting substitution model and the maximum likelihood phylogenetic tree with ultra-bootstrapping were implemented in IQTREE webserver ^19,20^. The resulting tree was subjugated to a root-to-tip regression analysis in TempEst^21^ to assess the clock-likeness regarding the data. A positive correlation was observed between sampling time and root-to-tip genetic divergence indicating the suitability of the dataset for molecular clock analysis using the Beast v1.10.4 package. The KHY+I substitution model with the uncorrelated lognormal relaxed clock, in addition to the coalescent exponential population growth model, were applied^22^. The MCMC chains were run for 200 million iterations and sampled every 10,000 cycles, or generations, with 10% discarded as burn in. We explored the effective sample sizes in Tracer (ESS>200)^23^. Moreover, to explore the phylogeographic diffusion of SARS-COV-2 in continuous space, the lognormal relaxed random walk diffusion model and a lognormal uncorrelated relaxed clock model were implemented in the same package, were next employed. Thus, the Maximum clade credibility tree was visualized in SpreaD3^24^.

Lineage assignment of the Hungarian sequences was performed using the PANGOLIN (Phylogenetic Assignment of Named Global Outbreak LINeages) software v1.0, which uses a recently published lineage nomenclature ^25,26^.

The datasets generated during and analysed during the current study are available in the NDEx-The Network Data Exchange repository, [http://www.ndexbio.org/#/network/2c66e15b-8eeb-11ea-aaef-0ac135e8bacf].

## Acknowledgements

On behalf of the project, “Genomic Epidemiology of SARS-CoV-2 in Hungary” we are grateful for the usage of MTA Cloud (https://cloud.mta.hu/) which significantly aided in achieving the results published in this paper. Gabor Kemenesi was supported by the Janos Bolyai Research Scholarship of the Hungarian Academy of Sciences. Balazs A Somogyi was supported by the ÚNKP-19-3 New National Excellence Program of the Ministry For Innovation and Technology. The research was performed in collaboration with the Genomics and Bioinformatics Core Facility at the Szentágothai Research Center of the University of Pécs. Bioinformatics infrastructure was supported by ELIXIR Hungary (http://elixir-hungary.org/).

## Authors Contribution

Conceptualization: GK, SZ, BAS, KB; Sample processing: GET, PU, ÁN, CIP, GCP, GD, GT; Sequencing: GET, PU; Bioinformatic support: RH, AGY; Sequence manipulation and analysis: SZ, BAS, KB, NS, PMS, IS, AB, RL, ST; Writing and editing: GK, BAS, SZ, FJ; Medical and epidemiological revision: VZ, JK; Supervising: FJ, GK. All authors reviewed the manuscript.

---

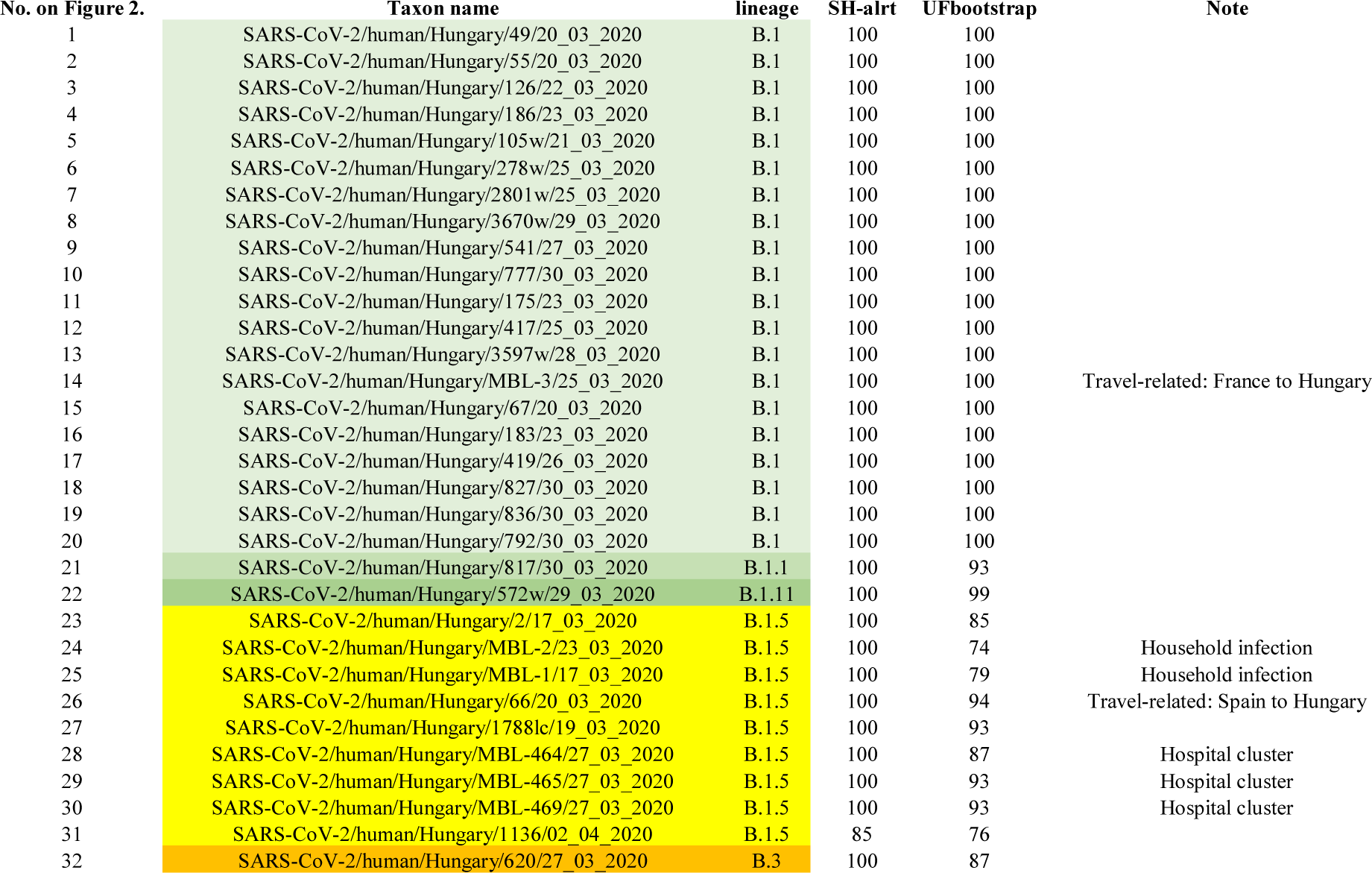

## Supplementary Data

**Supplemetary Figure 1.**
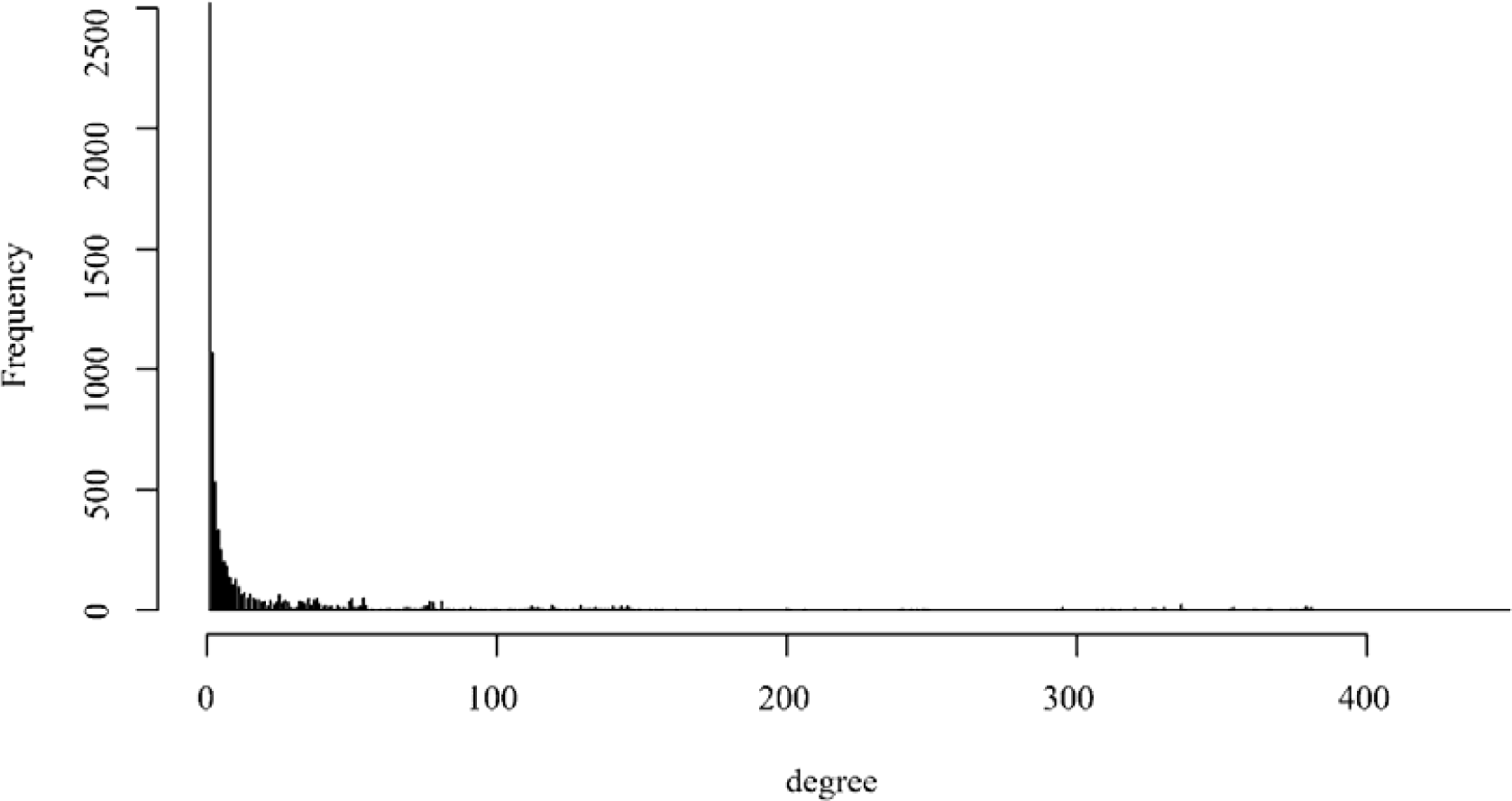
Degree distribution representing the haplotype network analysis

**Supplementary Fig. 2.**
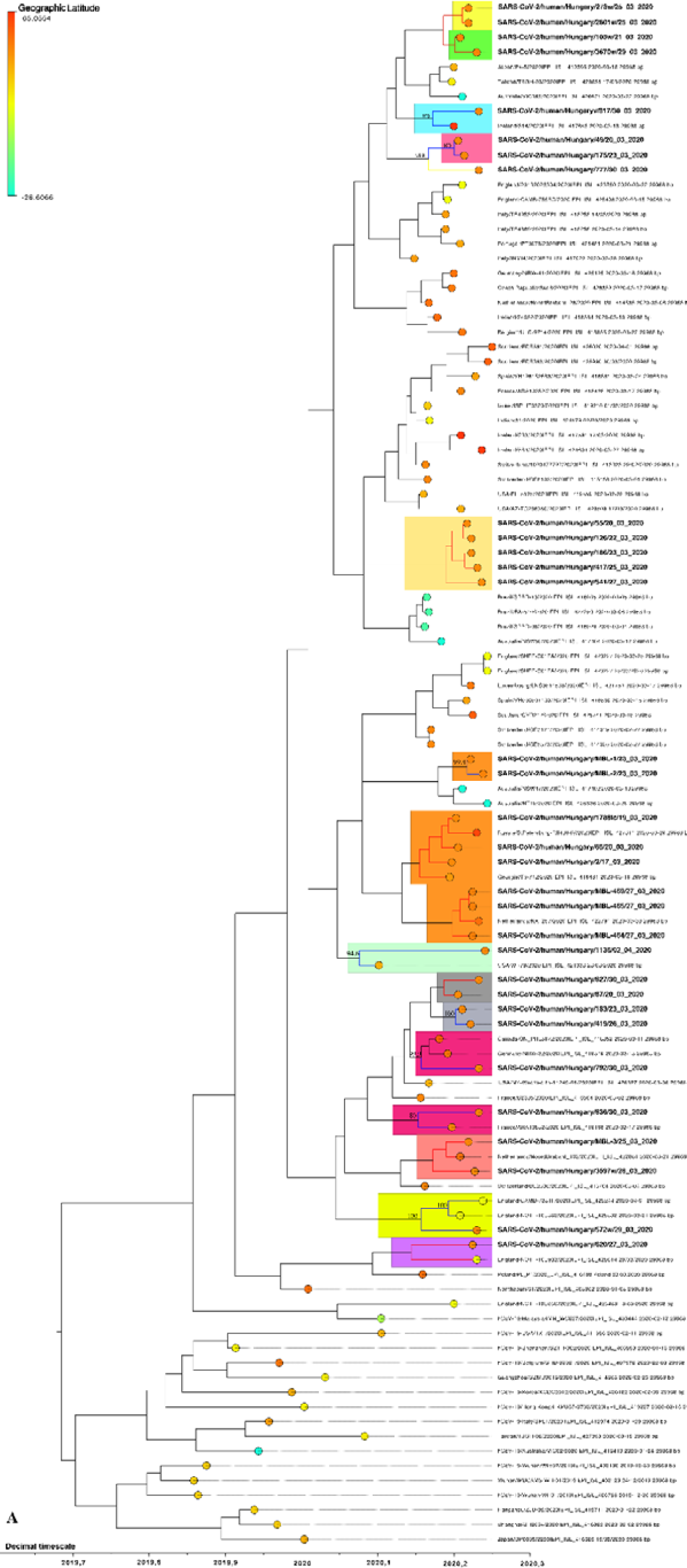
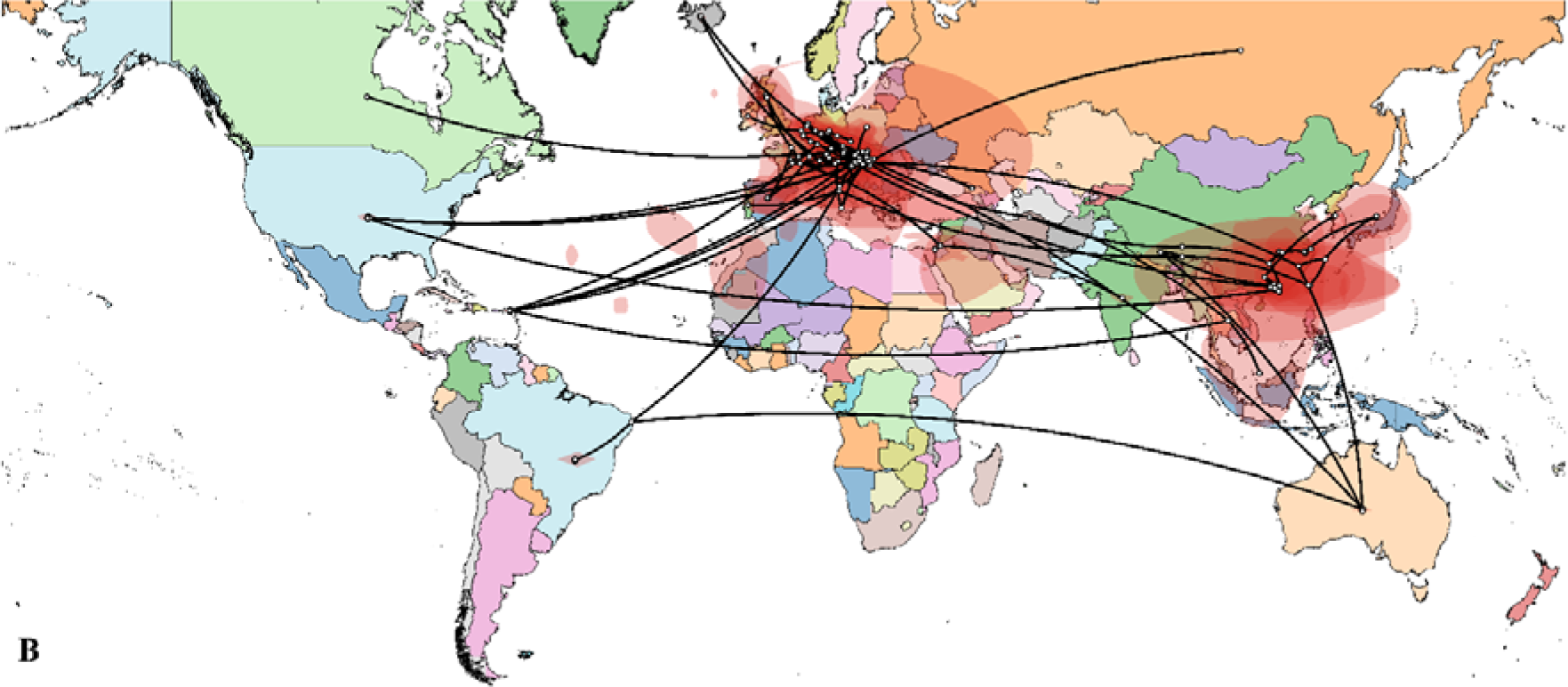
**A:** Time calibrated phylogenetic and phylogeographic visualization of 105 complete SARS-CoV-2 genomes compared to the 32 Hungarian strains. Sequences of this study are highlighted in bold face, colored clades are representing Nextstrain analysis clustering. **B**: The map is a visualization of the sequences presented at the phylogenetic tree.

**Supplementary Fig. 3.**
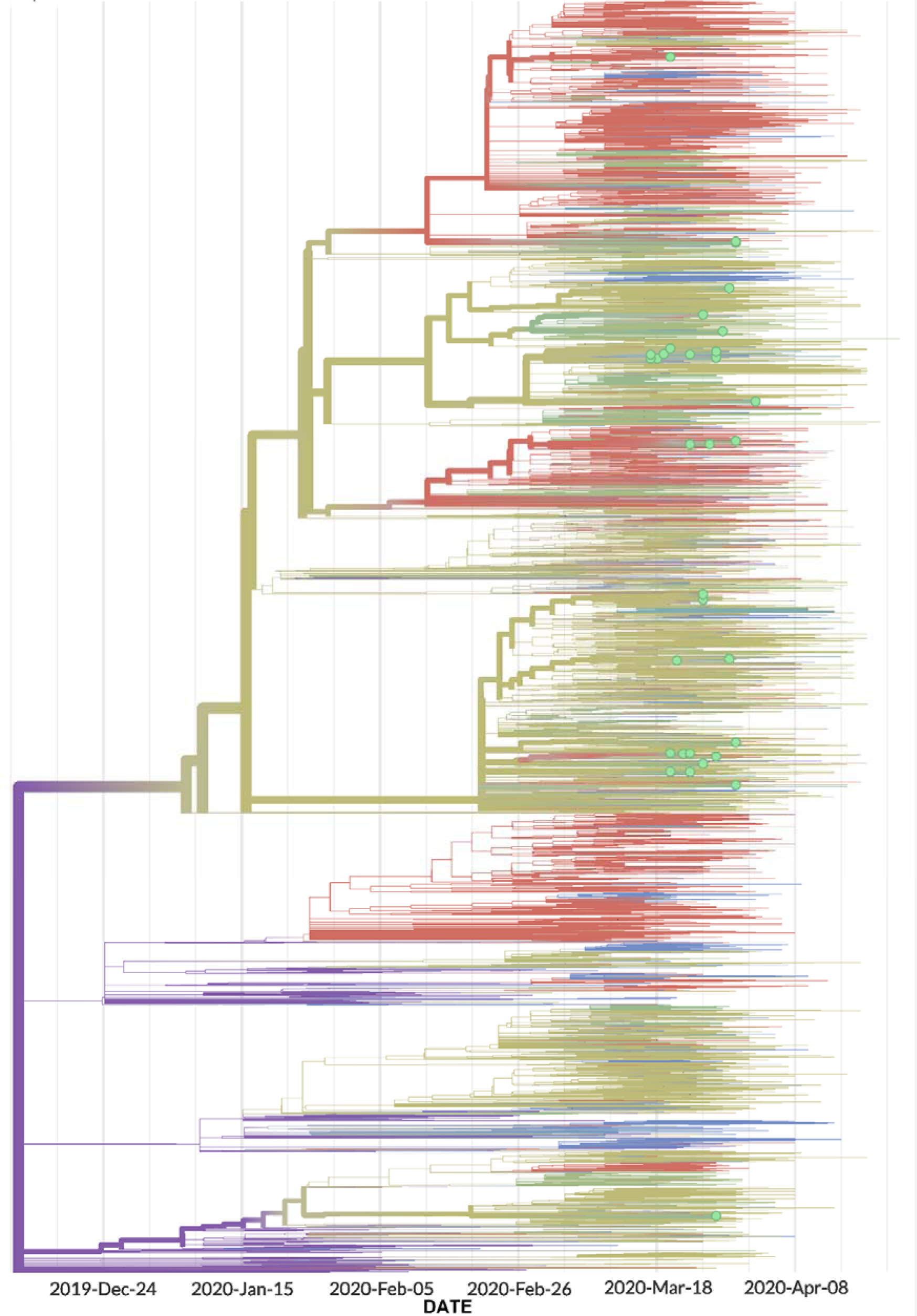
Visualization of Hungarian sequence dataset with Nextstrain local workflow. Showing 35 (32 from this manuscript and three additional from GISAID database) Hungarian sequences compared to 10,869 genomes sampled between Mar 2020 and Apr 2020.

